# MCM10 and RECQL4 have cooperative and redundant roles in activating the CMG helicase during the replication initiation

**DOI:** 10.64898/2026.04.01.715782

**Authors:** Atabek Bektash, Xiaoxuan Zhu, Yuki Hatoyama, Atsushi Toyoda, Masato T. Kanemaki

## Abstract

DNA replication initiation requires activation of the CMG helicase to establish the replisome. This process involves the extrusion of single-stranded DNA (ssDNA) from the central channel of MCM double hexamers, allowing the two CMG helicases to pass each other; however, the factors that mediate this process in human cells remain unclear. We show that degron-mediated depletion of either MCM10 or RECQL4 alone causes only mild replication defects, whereas simultaneous depletion of both proteins completely blocks CMG activation. ChIP-seq analyses demonstrate that RECQL4 localises to replication initiation zones (IZs) independently of MCM10, whereas MCM10 recruitment to IZs is enhanced upon RECQL4 depletion, suggesting RECQL4 primarily functions in CMG activation, and MCM10 acts as a backup or supporting factor. Rescue experiments further indicate that RECQL4 cooperates with MCM10 through direct interaction, and that their ssDNA-binding activity underlies their functional overlap. We propose MCM10 and RECQL4 act cooperatively and redundantly to promote CMG activation.

## Introduction

Faithful replication of genomic DNA is essential for cellular proliferation. In eukaryotic cells, DNA replication is tightly coupled to the cell cycle and occurs during S phase, in which tens of thousands of replication forks are generated. Replication initiation is the key regulatory step, temporally divided into two processes: licensing and firing (Costa and Diffley 2022; Zhu and Kanemaki 2024). Licensing occurs during G1 phase, when two MCM2–7 hexamers are recruited to DNA to form an MCM double hexamer (MCM-DH) through the action of the origin recognition complex (ORC), CDC6 and CDT1(**Fig. 1A, Licensing**). In this configuration, MCM2–7 is assembled to a head-to-head orientation and encircles double-stranded DNA (dsDNA), rendering it inactive as a replicative helicase. Firing occurs in S phase upon the activation of two kinases, DDK and S-CDK. The first step of firing is the recruitment of CDC45 and GINS to MCM-DH to form the replicative CDC45–MCM–GINS (CMG) helicase (**Fig. 1A, CMG assembly**). This step requires TRESLIN–MTBP, TOPBP1 and DONSON. The second step of firing is CMG activation, during which the two CMG helicases are separated and single-stranded DNA (ssDNA) is extruded from the central channel of the MCM ring. This allows each CMG helicase to encircle the leading-strand template and pass one another, thereby transitioning into two active replisomes (**Fig. 1A, CMG activation**). Importantly, the factors involved in CMG activation in human cells have remained unknown.

**Figure 1.**
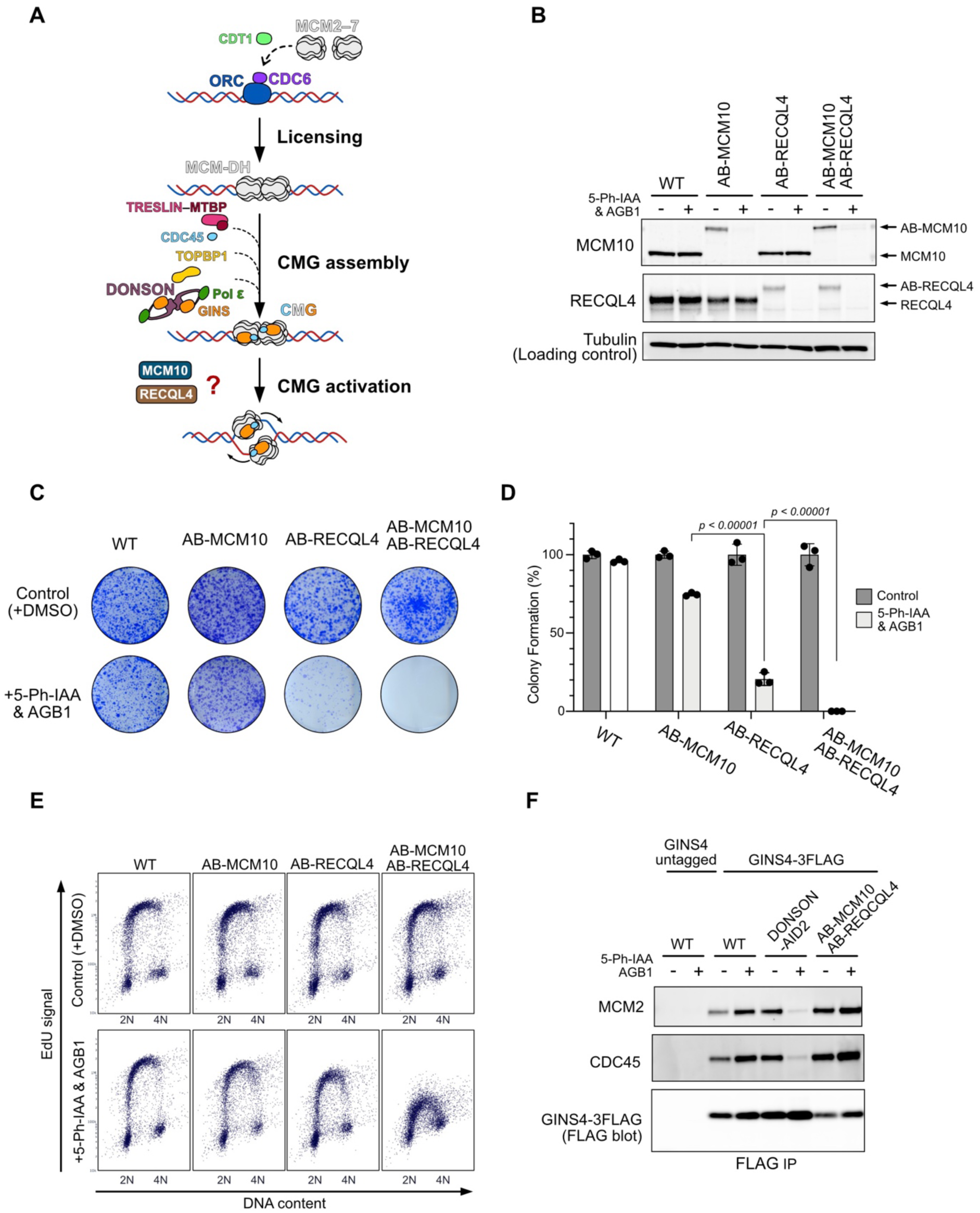
**(A)** Schematic representation of DNA replication initiation in human cells. **(B)** Immunoblot showing depletion of AB-MCM10, AB-RECQL4 or both proteins. Antibodies used for detection are indicated. The indicated cells were treated with 5-Ph-IAA and AGB1 for 6 h. **(C)** Representative images of stained colonies from the colony formation assay. The indicated cells were seeded at 1,000 cells per well and cultured for 7 days in the presence or absence of 5-Ph-IAA and AGB1. **(D)** Quantification of colony formation efficiency in the indicated cells. Data are presented as mean ± SD (*n* = 3 replicates; one-way ANOVA, planned pairwise comparison) and were normalised to the untreated cells. *p*-values were calculated for specific comparisons (AB-MCM10 vs AB-RECQL4; AB-RECQL4 vs AB-MCM10 and AB-RECQL4). **(E)** EdU incorporation and DAPI staining of the indicated cells treated with or without 5-Ph-IAA and AGB1 for 12 h. **(F)** Immunoblot analysis of CMG formation. GINS4-3×FLAG was immunoprecipitated from cells depleted of DONSON or MCM10/RECQL4. Antibodies used for detection are indicated.

Previously, we and other groups reported that Mcm10 is involved in CMG activation in yeast (Watase et al. 2012; Kanke et al. 2012; Deursen et al. 2012). More recently, biochemical and structural studies using yeast replication proteins have elucidated this process in detail (Douglas et al. 2018; Henrikus et al. 2024). MCM10 is conserved in humans and contains a conserved oligonucleotide/oligosaccharide-binding (OB)-fold domain with a high affinity for ssDNA (Warren et al. 2009; Eisenberg et al. 2009; Baxley and Bielinsky 2017). MCM10 is reported to be involved in replication initiation in human cells (Xu et al. 2009; Im et al. 2009; Izumi et al. 2017); however, its role in this process remains elusive.

RECQL4 is another enigmatic protein reported to be involved in replication initiation. The N-terminal region of RECQL4 shares homology with yeast replication factor Sld2, whereas the protein also has a long C-terminal extension harbouring a RECQ helicase domain (Chu and Hickson 2009; Croteau et al. 2012). Studies using *Xenopus* egg extracts demonstrated that the N-terminal region is essential for DNA replication, while the RECQ helicase domain is dispensable (Sangrithi et al. 2005; Matsuno et al. 2006). Consistently, chicken DT40 cells and mice expressing a RECQL4 mutant without the RECQ helicase domain are viable (Hoki et al. 2003; Mann et al. 2005; Abe et al. 2011). Interestingly, mutations within the RECQ helicase domain cause several autosomal recessive disorders, including Rothmund-Thomson syndrome (RTS), RAPADILINO syndrome (RAPA) and Baller-Gerold syndrome (BGS), which are associated with growth retardation, radial defects and cancer predisposition (Siitonen et al. 2009). Despite this, it remains unclear whether the RECQ helicase domain of RECQL4 has any contribution to replication initiation or instead functions in other DNA transactions (Croteau et al. 2012; 2014).

Based on the requirement of yeast Sld2 for CMG assembly (Muramatsu et al. 2010), the N-terminus of RECQL4 was thought to play a similar role in vertebrate cells. However, recent studies identified DONSON as fulfilling this role in vertebrate CMG assembly (Lim et al. 2023; Xia et al. 2023; Hashimoto et al. 2023), prompting a reassessment of the role of RECQL4 in replication initiation. Consistent with this view, RECQL4 is dispensable for DNA association of CDC45 and GINS, suggesting that CMG formation occurs independently of RECQL4 (Sanuki et al. 2015). Moreover, a recent study reported that RECQL4 functions during CMG activation in *Xenopus* egg extracts (Terui et al. 2024).

To study the role of MCM10 and RECQL4 in replication initiation in human cells, we generated conditional degron mutants for MCM10, RECQL4 or both proteins. We found that depletion of either MCM10 or RECQL4 alone causes mild defects in DNA replication, whereas simultaneous depletion of both proteins halts CMG activation, indicating functional redundancy between MCM10 and RECQL4. Using chromatin immunoprecipitation followed by sequencing (ChIP-seq), we show that RECQL4 localises to replication initiation zones (IZs) in early S phase irrespective of the presence or the absence of MCM10. In contrast, MCM10 accumulates at IZs upon RECQL4 depletion, suggesting MCM10 can function as a backup or supporting factor for RECQL4. Rescue experiments using various RECQL4 mutants revealed that RECQL4 cooperates through its interaction with MCM10, and their ssDNA binding activity underlies their functional redundancy. Based on these findings, we propose a model for how MCM10 and RECQL4 promote CMG activation.

## Results & Discussion

To study the role of MCM10 and RECQL4 in DNA replication initiation, we initially attempted to generate knockout cell lines in the human colorectal cell line HCT116. We aimed to knock-in a puromycin-resistant cassette into exon 3 of the *MCM10* gene using CRISPR–Cas9-mediated genome editing (**Fig. S1A**). Among the puromycin-resistant clones, genomic PCR analysis revealed that 16% harboured biallelic insertion of the selection cassette (**Fig. S1B**). However, subsequent Western blot analysis using an anti-MCM10 antibody showed that all of these biallelic clones expressed a truncated MCM10 protein that was not detected in the wild-type cells (**Fig. S1C**). We interpreted this smaller MCM10 product as being expressed from a downstream exon, likely representing a compensatory mechanism that suppresses complete loss of MCM10. These results suggest that the complete loss of MCM10 severely compromises cell viability or is lethal, consistent with a previous report (Lim et al. 2011; Baxley et al. 2021). Similarly, we attempted to knock-in a hygromycin-resistant cassette into exon 3 of the *RECQL4* gene (**Fig. S1D**). However, none of the hygromycin-resistant clones exhibited biallelic insertion, suggesting that the loss of RECQL4 is lethal in HCT116 cells (**Fig. S1E**). This observation contrasts with a recent report showing successful RECQL4 knockout in human osteosarcoma U2OS cells (Padayachy et al. 2024) and suggests that the requirement for RECQL4 may differ among cell types.

Because we were unable to generate knockout cell lines, we next turned to a conditional degron-based approach. We recently reported that combining an improved auxin-inducible degron (AID2) with a PROTAC-based BromoTag enhances target protein degradation (**Fig. S2A**) (Hatoyama et al. 2024). We therefore fused a tandem degron tag composed of mini-AID and BromoTag to the N-terminus of MCM10 and RECQL4, hereafter referred to as AB-MCM10 and AB-RECQL4, respectively. We confirmed the target protein depletion was rapid and efficient in the cells expressing AB-MCM10, AB-RECQL4 or both degron-tagged proteins (**Fig. 1B**). We then examined the effects of protein depletion on colony formation. As shown in **Fig. 1C, D**, depletion of either AB-MCM10 or AB-RECQL4 showed colony formation defects, with AB-RECQL4 depletion producing a more severe phenotype. Notably, simultaneous depletion of AB-MCM10 and AB-RECQL4 resulted in complete loss of viability.

We next examined the impact of target protein depletion on DNA synthesis during S phase (**Fig. 1E**). In the absence of degradation inducer, 5-Ph-IAA and AGB1, all degron-tagged cell lines exhibited EdU incorporation levels comparable to those of wild-type cells. In contrast, depletion of either AB-MCM10 or AB-RECQL4 resulted in reduced EdU incorporation, with a more pronounced defect in the AB-RECQL4 depleted cells. Importantly, simultaneous depletion of both AB-MCM10 and AB-RECQL4 caused a severe loss of EdU incorporation. The growth defect observed in **Fig. 1C** likely reflects the impaired DNA synthesis detected in **Fig. 1E**. Additionally, we confirmed that the depletion of both AB-MCM10 and AB-RECQL4 caused defective entry into S phase (**Fig. S2B**). Together, these results suggest that MCM10 and RECQL4 play partially redundant roles in DNA replication initiation.

A key event of replication initiation is the assembly of the CMG helicase. To determine whether MCM10 and RECQL4 function before or after CMG formation, we depleted both AB-MCM10 and AB-RECQL4 (**Fig. S2C**) and examined CMG assembly by immunoprecipitating 3×FLAG-tagged GINS4 (**Fig. 1F**). As a control, depletion of DONSON resulted in defective CMG formation consistent with a previous report (Lim et al. 2023). In contrast, in cells depleted of both AB-MCM10 and AB-RECQL4, CMG formation was comparable to that observed in wild-type cells, indicating that MCM10 and RECQL4 are not required for CMG assembly and suggesting they function during CMG activation (**Fig. 1A**).

If MCM10 and RECQL4 are involved in CMG activation, they are expected to colocalise with TRESLIN, which is required for CMG assembly and is localised at IZs. We therefore examined the genomic loci at which MCM10 and RECQL4 bind and their relationship to TRESLIN-binding loci. To this end, cells were synchronised in M phase using nocodazole and then released into the cell cycle, with samples collected at defined time points (**Fig. S3A, B**). We found that both MCM10 and RECQL4 accumulated on chromatin from early S phase (6 h after release) (**Fig. S3C**), consistent with the hypothesis that they function during CMG activation. We recently reported ligase-depletion OK-seq (LD-OK-seq) to map early-firing IZs, together with ChIP-seq analyses of MCM4 and TRESLIN (**Fig. 2A**) (Zhu et al. 2025). Following the same ChIP-seq protocol, we immunoprecipitated 3×FLAG-fused MCM10 or RECQL4, either in the presence or absence of the other protein. In the presence of RECQL4, MCM10 was detectable (**Fig. 2A, B, +RECQL4**). However, upon the depletion of RECQL4, MCM10 became clearly enriched at early-firing IZs, similar to the binding pattern of TRESLIN (**Fig. 2A, B, -RECQL4** and **Fig. S3D, E**). In contrast, RECQL4 was detected at comparable levels at early-firing IZs regardless of the presence or absence of MCM10 (**Fig. 2A, C**), and its location was similar to TRESLIN (**Fig. S3D, F**). We compared MCM10 and RECQL4 ChIP-seq profiles with IZs detected by LD-OK-seq and with TRESLIN ChIP-seq data (**Fig. 2D**). Among all IZs, early-firing IZs are marked by TRESLIN (note that ChIP-seq was carried out using early S phase cells) (Zhu et al. 2025). Regardless of the presence or absence of MCM10, RECQL4 exhibited a distribution pattern similar to that of TRESLIN. In contrast, MCM10 displayed a TRESLIN-like distribution only in the absence of RECQL4. The weak enrichment of MCM10 at early-firing IZs in the presence of RECQL4 is likely attributable to its broader distribution across the non–early-firing IZs (**Fig. 2D, +RECQL4**). Taken together, we concluded that RECQL4 plays a primary role during CMG activation, whereas MCM10 functions as a backup or supporting factor. This interpretation is consistent with the more severe defects in cell growth and DNA synthesis observed upon RECQL4 depletion compared with MCM10 depletion (**Fig. 1C–E**).

**Figure 2.**
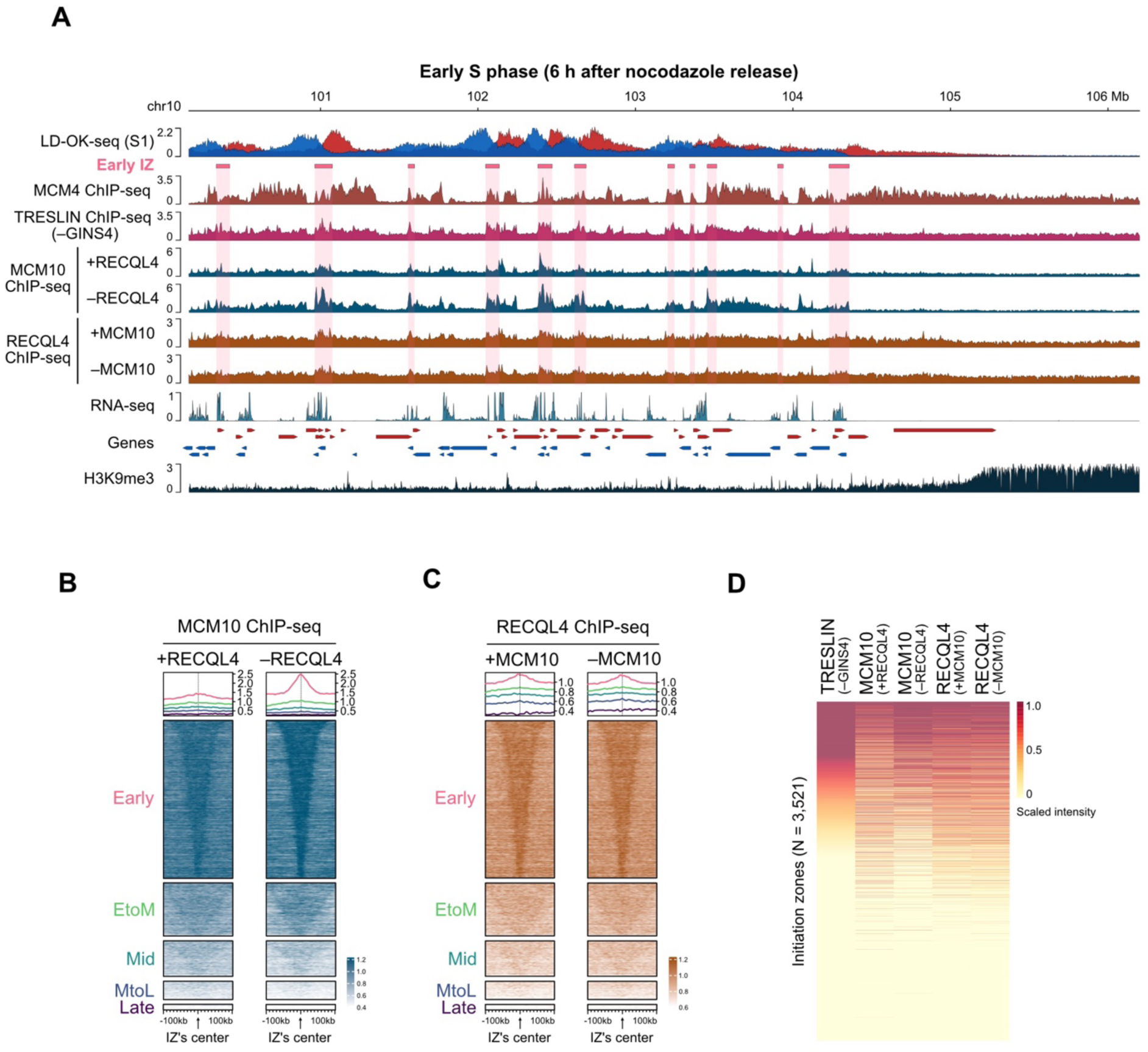
**(A)** Representative genomic profiles of LD-OK-seq (S1), ChIP-seq for MCM4 (G1), TRESLIN (S1), MCM10 (S1), and RECQL4 (S1), RNA-seq, and H3K9me3. Okazaki fragment reads from the Crick and Watson strands are shown in red and blue, respectively. Early-firing IZs are highlighted in pink. All ChIP-seq data are presented as input-normalised signal ratios. TRESLIN ChIP-seq profiles are obtained from GINS4-depleted cells (Zhu et al. 2025). RNA-seq data are displayed as log2-transformed TPM values. Genes are annotated with coloured arrows indicating transcriptional directions. **(B)** Heatmaps showing MCM10 enrichment at IZs in the presence or absence of RECQL4. IZs are categorised according to their activation timing in S phase. IZs are centered with ±100 kb flanking regions, and ordered by size. **(C)** Heatmaps showing RECQL4 enrichment at IZs in the presence or absence of MCM10. IZs are categorised according to their activation timing in S phase. IZs are centered with ±100 kb flanking regions, and ordered by size. **(D)** Relationship between TRESLIN, MCM10 and RECQL4 accumulation at IZs under the indicated conditions. All ChIP-seq data were obtained from cells in early S phase (S1).

Based on our finding that RECQL4 plays a primary role in CMG activation, we next focused on RECQL4. To this end, we rescued HCT116 cells expressing AB-RECQL4 by introducing RECQL4 mutants (**Fig. 3A** and **S4A)**. The K508A mutant carries a K-to-A substitution within the Walker A motif that abolishes ATPase activity (Rossi et al. 2010). The N1 mutant is a truncation lacking the RECQ helicase family domain. The N2 mutant is a shorter truncation that retains the Sld2-like and MCM10-binding domains (Sangrithi et al. 2005; Xu et al. 2009; Kliszczak et al. 2015). All mutants were fused to an SV40 nuclear localisation signal (NLS), and their nuclear localisation was confirmed (**Fig. S4B**). The AB-RECQL4 parental cells showed a defect in colony formation upon depletion of AB-RECQL4 (**Fig. 3B, C**). Expression of RECQL4 wild-type and all RECQL4 mutants restored colony formation to similar levels, suggesting that all mutants are capable of supporting CMG activation in the presence of MCM10. These results are consistent with the reports showing that the RECQ helicase domain of RECQL4 is dispensable for viability in cultured cell lines and mice (Hoki et al. 2003; Mann et al. 2005; Abe et al. 2011). In contrast, the AB-MCM10 AB-RECQL4 cells failed to form colonies when both proteins were depleted (**Fig. 3D, E**). This lethality was rescued by RECQL4 wild-type, K508A and N1 to similar extents, but not by N2. These results indicate that the N2 fragment supports CMG activation only in the presence of MCM10.

**Figure 3.**
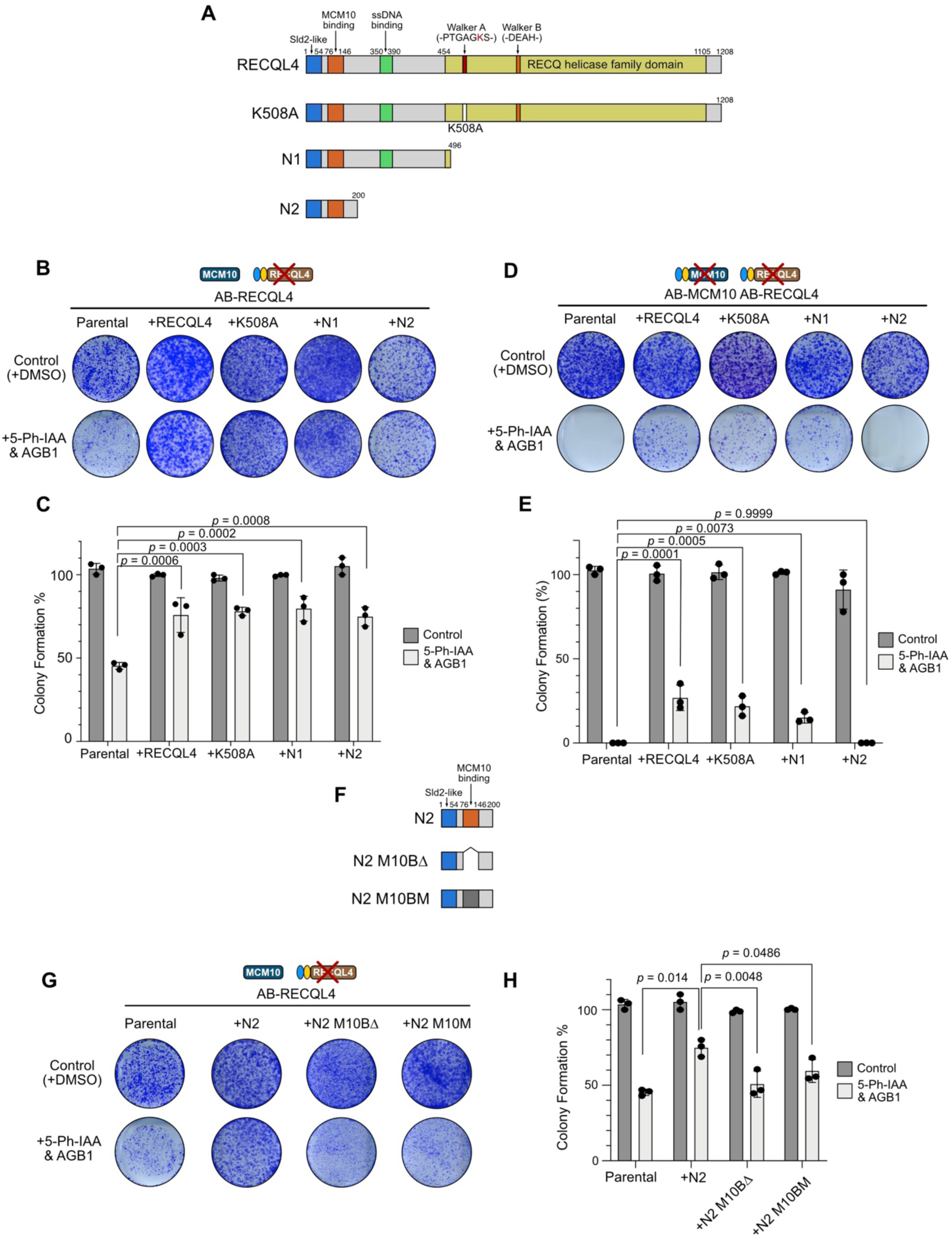
**(A)** Schematic illustration of RECQL4, showing its domains and the RECQL4 mutants used in the study. **(B, D)** Representative images of stained colonies. The indicated cells were seeded at 1,000 cells per well and cultured for 7 days in the presence or absence of 5-Ph-IAA and AGB1. **(C, E)** Quantification of colony formation efficiency in the indicated cells. Data are presented as mean ± SD (*n* = 3 replicates; one-way ANOVA, multiple comparison) and were normalised to the untreated cells. *p*-values were calculated in comparison to the parental cell line. **(F)** Schematic illustration of RECQL4-N2 and two mutants deficient in interaction with MCM10. **(G)** Representative images of stained colonies. Cells expressing RECQL4-N2 or the indicated mutants were seeded at 1,000 cells per well and cultured for 7 days in the presence or absence of 5-Ph-IAA and AGB1. **(H)** Quantification of colony formation efficiency of cells expressing the indicated RECQL4-N2 mutant. Data are presented as mean ± SD (*n* = 3 replicates; one-way ANOVA, multiple comparison) and were normalised to the untreated cells. *p*-values were calculated in comparison to the N2 expressing cell line.

RECQL4 is reported to interact with MCM10 (Xu et al. 2009), and we confirmed this interaction (**Fig. S4C**). The RECQL4 N2 mutant contains an MCM10-binding domain (**Fig. 3A**) (Kliszczak et al. 2015), which is predicted to interact with the OB-fold domain of MCM10 (**Fig. S4D**). We hypothesised that the N2 mutant functions by supporting MCM10-mediated CMG activation through its interaction with MCM10. To test this, we introduced either a deletion (N2-M10BΔ) or point mutations (N2-M10M) to the N2 fragment (**Fig. 3F** and **S4E**) and expressed these mutants in the AB-RECQL4 cells (**Fig. S4A, B**). Neither N2-M10BΔ nor N2-M10M restored colony formation, despite the presence of endogenous MCM10 (**Fig. 3G, H**). These results are consistent with the view that RECQL4 can function cooperatively with MCM10 during CMG activation.

An interesting phenotypic difference between the RECQL4 N1 and N2 mutants is that N1 functions in the absence of MCM10, but N2 does not (**Fig. 3B–E**). The N1 mutant contains an ssDNA-binding domain (Keller et al. 2014; Sedlackova et al. 2015) (**Fig. 4A** and **S5A**). Similarly, the OB-fold domain in MCM10 is known to interact with ssDNA (**Fig. 4A** and **S5B**) (Robertson et al. 2008; Warren et al. 2008). We hypothesised that the N1 mutant supports viability in the absence of MCM10 because it is capable of binding ssDNA. To test this hypothesis, we introduced point mutations that disrupt ssDNA binding in RECQL4 (**Fig. 4A** and **S5C**) and examined whether these RECQL4-DBM and N1-DBM mutants support viability. We found that neither RECQL4-DBM nor N1-DBM supported cell viability in the absence of endogenous MCM10 (**Fig. 4B, C**), whereas both restored colony formation in the presence of MCM10 (**Fig. S5D, E**). These results are consistent with the view that ssDNA binding by RECQL4 and MCM10 is critical for CMG activation, and this activity must be provided by at least one of the two proteins (**Fig. 4D**).

**Figure 4.**
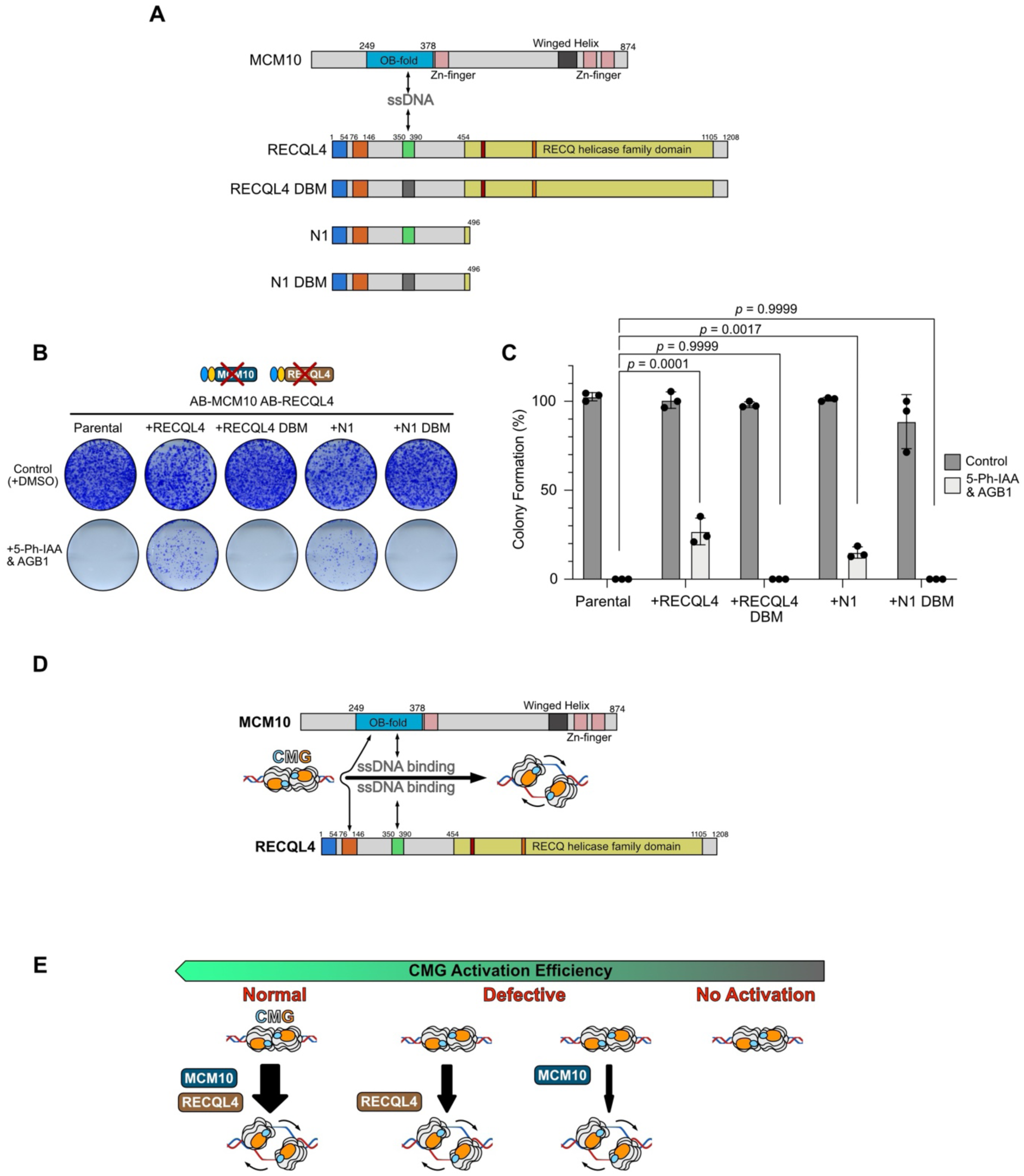
**(A)** Schematic illustration of MCM10 and RECQL4, showing their domains and the RECQL4 mutants deficient in ssDNA interaction. **(B)** Representative images of stained colonies. The indicated cells were seeded at 1,000 cells per well and cultured for 7 days in the presence or absence of 5-Ph-IAA and AGB1. **(C)** Quantification of colony formation efficiency in the indicated cells. Data are presented as mean ± SD (*n* = 3 replicates; one-way ANOVA, multiple comparison) and were normalised to the untreated cells. *p*-values were calculated in comparison to the parental cell line. **(D)** Graphic summary illustrating the domains of MCM10 and RECQL4 and their role in promoting CMG activation. **(E)** Graphic summary illustrating the functional redundancy between MCM10 and RECQL4 and its relationship to the CMG activation efficiency.

Taken together, we propose that both MCM10 and RECQL4 contribute to CMG activation (**Fig. 4E**), with RECQL4 playing a primary role in this process. This conclusion is supported by colony formation assay showing that RECQL4 depletion has a severe impact on both colony formation and EdU incorporation (**Fig. 1**), as well as by the observation that RECQL4 localises to early-firing IZs regardless of the presence of MCM10 (**Fig. 2**). The potential direct interaction between the Sld2-like domain of RECQL4 and the N-terminus of MCM2 may contribute to RECQL4 loading onto CMG helicase (**Fig. S5F**). Our results indicate that RECQL4 can promote CMG activation even without MCM10 (**Fig. 4E**), whereas MCM10 appears to function as a backup or supporting factor in this process. Nevertheless, we also propose that RECQL4 and MCM10 can act cooperatively during CMG activation, as the MCM10-binding region in RECQL4-N1 is required for efficient colony formation (**Fig. 3F–H**).

This interpretation is consistent with previous reports demonstrating that both MCM10 and RECQL4 are required for DNA replication initiation (Xu et al. 2009; Im et al. 2009; Kliszczak et al. 2015; Im et al. 2015). In the absence of RECQL4, MCM10 accumulates at early-firing IZs (**Fig. 2**). This observation suggest that, although MCM10 can promote CMG activation in the absence of RECQL4, it does so less efficiently, possibly reflecting prolonged residence of MCM10 on the CMG helicase (**Fig. 4E**). Recruitment of MCM10 to the assembled CMG may involve its reported MCM2–7 binding domain (Izumi et al. 2004) in addition to its interaction with RECQL4 (**Fig. 3F–H** and **S4C**). Although either MCM10 or RECQL4 can activate CMG helicase, it is important to note that both MCM10 and RECQL4 are required for efficient CMG activation to fully support normal cell proliferation (**Fig. 1C, D**). This requirement likely explains why we were unable to generate a knockout cell line for either MCM10 or RECQL4 (**Fig. S1**).

Our findings further indicate that functional redundancy between MCM10 and RECQL4 is conferred by their ability to bind ssDNA (**Fig. 4A–C** and **Fig. S5A–E**). During CMG activation, ssDNA must be extruded from the MCM hexameric ring to allow two CMG helicases to pass over each other. We hypothesise that, following the conformational change in the MCM-DH induced by DONSON (Cvetkovic et al. 2023), MCM10 and RECQL4 cooperate to promote ssDNA extrusion, thereby enabling the CMG helicase to encircle the leading-strand template DNA. Interestingly, AlphaFold3 predictions suggest that when RECQL4-N1 binds to the N-terminus of MCM2, its ssDNA binding domain is positioned close to the DNA between MCM-DH, as if positioned to capture extruded ssDNA (**Fig. S5F**). Additionally, the MCM10-interacting domain of RECQL4 is located close by. When both MCM10 and RECQL4 are depleted, this process fails, resulting in defective CMG helicase activation (**Fig. 4E**).

Our conclusion that RECQL4 promotes CMG activation is consistent with a recent observation in *Xenopus* egg extracts, which reported that RECQL4 functions after CMG formation (Terui et al. 2024). In contrast, Thakur et al. recently reported that RECQL4 suppresses TRESLIN–MTBP loading at dormant origins, thereby preventing CMG formation (Thakur et al. 2025). We compared our IZ and RECQL4 ChIP-seq data obtained from HCT116 cells in early S phase with the RECQL4 ChIP-seq data from asynchronous HCT116 cells reported by Thakur et al. (**Fig. S6**). We found that the RECQL4 ChIP-seq peaks reported by Thakur et al. neither colocalise well with IZs (**Fig. S6A, B**) nor align with our RECQL4 and TRESLIN ChIP-seq profiles (**Fig. S6C**). The reason for this discrepancy is currently unclear and needs to be clarified in the future.

RECQL4 contains an N-terminal Sld2-like domain and a C-terminal conserved RECQ helicase domain (**Fig. 3A**). Yeast Sld2 is required for CMG formation (Muramatsu et al. 2010), and in addition to this function, Sld2 may also be involved in CMG activation in yeast, similar to RECQL4 in humans. Mutations within the RECQ helicase domain of RECQL4 are associated with human genetic disorders, RTS, RAPA and BGS (Siitonen et al. 2009; Chu and Hickson 2009; Croteau et al. 2012; 2014). In the present study, we were unable to define a clear role for the RECQ helicase domain in CMG activation. It remains possible that this domain contributes to CMG activation in a manner that is subtle or context-dependent and therefore not readily detectable by our experimental assays. Alternatively, the RECQ helicase domain of RECQL4 may primarily function in DNA repair-related processes rather than replication initiation. Consistent with this idea, RECQL4 has been implicated in multiple DNA transactions, including base-excision repair and double-strand break repair (Schurman et al. 2009; Singh et al. 2010; Lu et al. 2017). Thus, while the N-terminal functions of RECQL4 appear to play a dominant role in CMG activation, the conserved helicase domain may mediate replication-independent functions that underlie the disease phenotypes associated with RECQL4 mutations.

In summary, we demonstrated that MCM10 and RECQL4 contribute to CMG activation in a cooperative and redundant manner. Using single-molecule imaging with *Xenopus* egg extracts, a similar conclusion was recently reported in a preprint study (Terui et al. 2025). Defining the precise molecular roles of MCM10 and RECQL4 in CMG activation will require biochemical and structural analyses of initiation intermediate complexes purified from replicating chromatin or reconstituted in vitro using human proteins.

## Materials & Methods

### Cell culture and cell cycle synchronisation

HCT116 cells were cultured in McCoy’s 5A, supplemented with 10% FBS, 2 mM L-glutamine, 100 U/ml penicillin and 100 µg/mL streptomycin at 37 °C in a 5% CO_2_ incubator. To synchronise cells, nocodazole (Sigma, #M1404) was added to the culture medium at a final concentration of 50 ng/ml. After 14-16 h, the nocodazole-containing media was removed, and the cells were washed twice with fresh media. Following the nocodazole release, 1 µM 5-Ph-IAA (AID2 degradation inducer) and 0.5 µM AGB1 (BromoTag degradation inducer) were added for the desired period of time. 5-Ph-IAA was synthesised as previously described (Yesbolatova et al. 2020). AGB1 was commercially obtained (Tocris #7686).

### Colony formation assay

In a 6-well plate, 5000 cells were seeded and cultured in the presence or absence of the indicated ligands for 7 days. The culture medium was exchanged with fresh medium containing appropriate ligands on day 4. Cells were fixed and stained with a crystal violet solution (6.0% glutaraldehyde and 0.5% crystal violet). Colonies were quantified using ImageJ (Fiji) and normalised to the mean untreated value of the same cell line, set to 100%. To account for intrinsic differences in baseline growth potential between cell lines, 5-Ph-IAA- and AGB1-treated values were normalised to the mean control value (DMSO-treated condition) of the corresponding cell line.

Statistical analyses were performed on the normalised values obtained from 5-Ph-IAA- and AGB1 conditions. Differences among cell lines were assessed using one-way analysis of variance (ANOVA) with a pooled variance model. Comparisons between each cell line and the reference cell line (parental, otherwise specified) were conducted using Dunnett’s multiple-comparisons test. Additional comparisons between specified cell lines were performed using planned pairwise comparisons with the pooled variance from the ANOVA. All experiments were performed with three biological replicates.

### Plasmids

All plasmids used in this study are listed in **Table S1**. PiggyBac plasmids used to rescue RECQL4 depletion encode either wild-type or mutant RECQL4 as a fusion protein with a 3×FLAG tag and an SV40 NLS.

### Cell lines

All cell lines used in this study are listed in **Table S2**. HCT116 cell lines expressing degron-fused protein(s) were generated following the protocol previously published (Saito and Kanemaki 2021; Hatoyama et al. 2024). All guide-RNA sequences used in this study are shown in **Table S3**. A piggyBac plasmid encoding RECQL4 mutants was transfected with pCMV-hyPBase encoding piggyBac transposase (Yusa et al. 2011). Subsequently, stable cells were selected using puromycin or blasticidin S.

### Protein detection

Cells were harvested by trypsin and washed with culture medium and PBS. Cells were lysed with RIPA buffer (25 mM Tris-HCl pH 7.5, 150 mM NaCl, 1% NP-40, 1% sodium deoxycholate, 0.1% SDS) containing complete protease inhibitor cocktail (Roche, #1187580001) for 30 min on ice. Subsequently, the tubes were centrifuged for 15 min at 4 °C, and then the supernatant was mixed with the same amount of SDS-sample buffer (Cosmo Bio, #423420) before heating at 95 °C for 5 min. The denatured protein samples were separated on a 7.5% or 10% TGX Stain-Free gel (BioRad) and transferred to a nitrocellulose membrane (Cytiva, #10600003). The membrane was processed with 1% skim milk in TBS-T for 15 min and incubated with primary antibody at 4 °C overnight. After washing with TBS-T, the membrane was incubated with corresponding secondary antibody for 1 h at RT. Proteins were detected by ChemiDoc Touch MP imaging system (BioRad), and the detected signals were quantified using Image Lab software (ver. 6.0.1, BioRad).

### Antibodies

All antibodies used in this study are listed in **Table S3**. **EdU labelling and quantification** For detecting DNA replication by EdU incorporation, cells were cultured with 10 µM EdU for 30 min before fixation. Incorporated EdU was stained using the Click-iT EdU Imaging Kit (Thermo Fisher Scientific, #C10339) according to the manufacturer’s instructions, and nuclear DNA was stained with DAPI. The EdU and DAPI signals were captured with an ECLIPSE Ti2 wide-field microscope (Nikon) through a CFI Plan Apo 10×/0.45 objective lens (Nikon) equipped with ORCA-FusionBT C15440 (Hamamatsu). Obtained images were processed using Fiji (ImageJ2, ver. 2.14.0) to perform nuclear segmentation. Background was removed using rolling-ball subtraction, after which EdU and DAPI signals were quantified. Scatter plots were generated using Python (Plotly Dash).

### Flow Cytometry

Trypsinised cells were washed once with a fresh medium and resuspended in 0.15 ml of ice-cold PBS. Cells were then fixed by adding 0.35 ml of ice-cold 100% ethanol dropwise while gently shaking. Fixed cells were stored at −20 °C overnight. After fixation, cells were washed once with PBS containing 1% BSA and resuspended in staining buffer (PBS, 1% BSA, 40 µg/ml propidium iodide, 50 µg/ml RNase A). Staining was performed at 37 °C for 30 min. Flow cytometry was performed using a BD Accuri C6 Cytometer.

### Separation of soluble and chromatin-bound proteins

To separate soluble and chromatin-bound proteins, cells were collected by trypsinisation and washed twice with PBS. Subsequently, cells were resuspended in EXB buffer (25 mM Tris-HCl pH 7.5, 150 mM NaCl) containing 0.2% Triton X-100, and then pelleted by centrifugation at 845 × g at 4 °C for 2 min. The supernatant was collected as the soluble protein fraction. The cell pellet was washed once with EXB containing 0.2% Triton X-100. The chromatin-bound protein was extracted by adding RIPA buffer to the pellet. After incubation on ice for 30 min, the sample was centrifuged at 18,400 × g at 4 °C for 10 min. The supernatant (chromatin-bound protein fraction) was collected. An equal volume of 2×SDS sample buffer was added to each sample, which was subsequently boiled at 95 °C for 5 min to denature proteins.

### Immunoprecipitation

Immunoprecipitation was conducted following established protocols (Zhu et al. 2025). Briefly, 2 × 10^7^ cells were collected and resuspended in 1 ml of IP lysis buffer (20 mM HEPES pH 7.5, 150 mM NaCl, 1 mM EDTA, 0.5% NP-40, 5% glycerol, 3 mM MgCl_2_). The buffer was supplemented with 100 U benzonase (Millipore, #70746) along with protease and phosphatase inhibitor cocktails (Roche, #11873580001 and #490684500). After 30-minute incubation at 10 °C, the lysate was clarified via two rounds of centrifugation (18,400 × g, 15 min, 4 °C). To minimize non-specific binding, the supernatant was pre-cleared using 0.75 mg Protein G Dynabeads (Invitrogen, #10004D) for 1 hour at 4 °C. This pre-cleared lysate was then incubated for overnight at 4 °C with anti-FLAG M2 antibody (Sigma, #F1804) conjugated to 1.5 µg Protein G Dynabeads. Following three washes with 1 ml of IP wash buffer, the protein complexes were eluted by boiling in 1×SDS sample buffer at 95 °C for 5 min.

### Chromatin immunoprecipitation sequencing (ChIP-seq)

ChIP-seq analysis was performed according to previously established methods (Zhu et al. 2025). Approximately 2 × 10^7^ cells were trypsinised, washed twice with 10 ml PBS, and resuspended in 1 ml PBS for crosslinking. This was achieved by adding an equal volume of fresh 2% paraformaldehyde (10 min, room temperature), followed by quenching with 125 mM glycine for 5 min. After sequential washes with PBS, PBS/0.5% NP-40, and PBS/10% glycerol, cell pellets were either flash-frozen in liquid nitrogen for storage at -80 °C or lysed immediately. For lysis, cells were incubated on ice for 10 min in 1 ml ChIP lysis buffer (25 mM HEPES pH 7.5, 140 mM NaCl, 1 mM EDTA, 0.5 mM EGTA, 0.5% Sarkosyl, 0.1% sodium deoxycholate, 0.5% Triton X-100, and Roche protease inhibitors). Chromatin fragmentation was performed using a Covaris S2 focused-ultrasonicator (5% duty factor, 100 cycles/burst, intensity 3) for 10 min. Following centrifugation (18,400 x g, 15 min, 4 °C), the supernatant was pre-cleared with 1.5 µg Protein G Dynabeads (Invitrogen, #10004D) for 1–2 h. A 2% input aliquot (20 µl) was reserved, while the remaining supernatant underwent immunoprecipitation overnight at 4 °C with 5 µg anti-FLAG M2 antibody (Sigma, #F1804) bound to Protein G Dynabeads. The beads were washed stringently with RIPA buffer, RIPA containing 300 mM NaCl, RIPA with 250 mM LiCl, and twice with TE buffer. DNA was eluted twice at 65 °C in TE + 1% SDS, and crosslinks were reversed at 65 °C for 4 h following RNase A and Proteinase K treatment. Finally, DNA was purified (Takara NucleoSpin), quantified (PicoGreen), and libraries were generated using the NEBNext Ultra II kit for 50 bp paired-end sequencing on an Illumina NovaSeq 6000.

### ChIP-seq data processing

Initial assessment of raw FASTQ sequencing reads was performed using FastQC (v0.11.9). We utilized Trim Galore (v0.6.6) for adapter removal (Illumina sequence: AGATCGGAAGAGC) via the --illumina flag. The processed reads were mapped to the GRCh38 (NCBI) reference genome with Bowtie2 (v2.2.5), employing stringent alignment criteria (--end-to-end --very-sensitive --no-mixed --no-discordant -I 0 -X 1000). Following alignment, SAMtools (v1.12) was used to convert outputs to BAM format and enforce quality filters, including a MAPQ score ≥ 20 and a requirement for proper pairing. PCR duplicates were identified and discarded using the fixmate and markdup suites. Genomic signal tracks were generated with MACS2 (v2.2.7.1) and converted to bigWig files. Final normalization was achieved through deepTools (v3.5.5) bigwigCompare, which calculated the ChIP/Input ratio across 100 bp genomic windows (--operation ratio --pseudocount 0 -bs 100). This pipeline was applied consistently to both newly generated and reprocessed public datasets.

## Data availability

Data generated in this study have been deposited in the Gene Expression Omnibus (GEO) database under the accession number GSE317126. Published datasets used in this study includes: Early S phase LD-OK-seq data in HCT116 GSE293198 [https://www.ncbi.nlm.nih.gov/geo/query/acc.cgi?acc=GSE293198]; MCM4 and TRESLIN ChIP-seq data in HCT116 GSE293195 [https://www.ncbi.nlm.nih.gov/geo/query/acc.cgi?acc=GSE293195]; RNA-seq data GSE293200 [https://www.ncbi.nlm.nih.gov/geo/query/acc.cgi?acc=GSE293200]; H3K9me3 ChIP-seq ENCSR000FCP [https://www.encodeproject.org/experiments/ENCSR000FCP/]; and RecQL4 ChIP-seq from Thakur et al. GSE276856 [https://www.ncbi.nlm.nih.gov/geo/query/acc.cgi?acc=GSE276856].

## Acknowledgements

We thank all members of the Kanemaki laboratory for discussion and support. AB is a MEXT Scholarship Fellow. We thank the NIG supercomputer facility for its support in data analysis. This work was supported by JSPS KAKENHI grants (JP21H04719, JP23H04925, JP25H00979 and JP22H04925 (PAGS)), JST CREST grant (JPMJCR21E6) to MTK.

## Author Contributions

This project was designed by AB and MTK with the help of the other authors. AB, XZ, and YH performed experiments. AT performed high-throughput sequencing. AB and MTK wrote the manuscript with contributions from the other authors.

