## Supplementary material for "MCM10 and RECQL4 have cooperative and redundant roles in activating the CMG helicase during the replication initiation": Figures S1-6

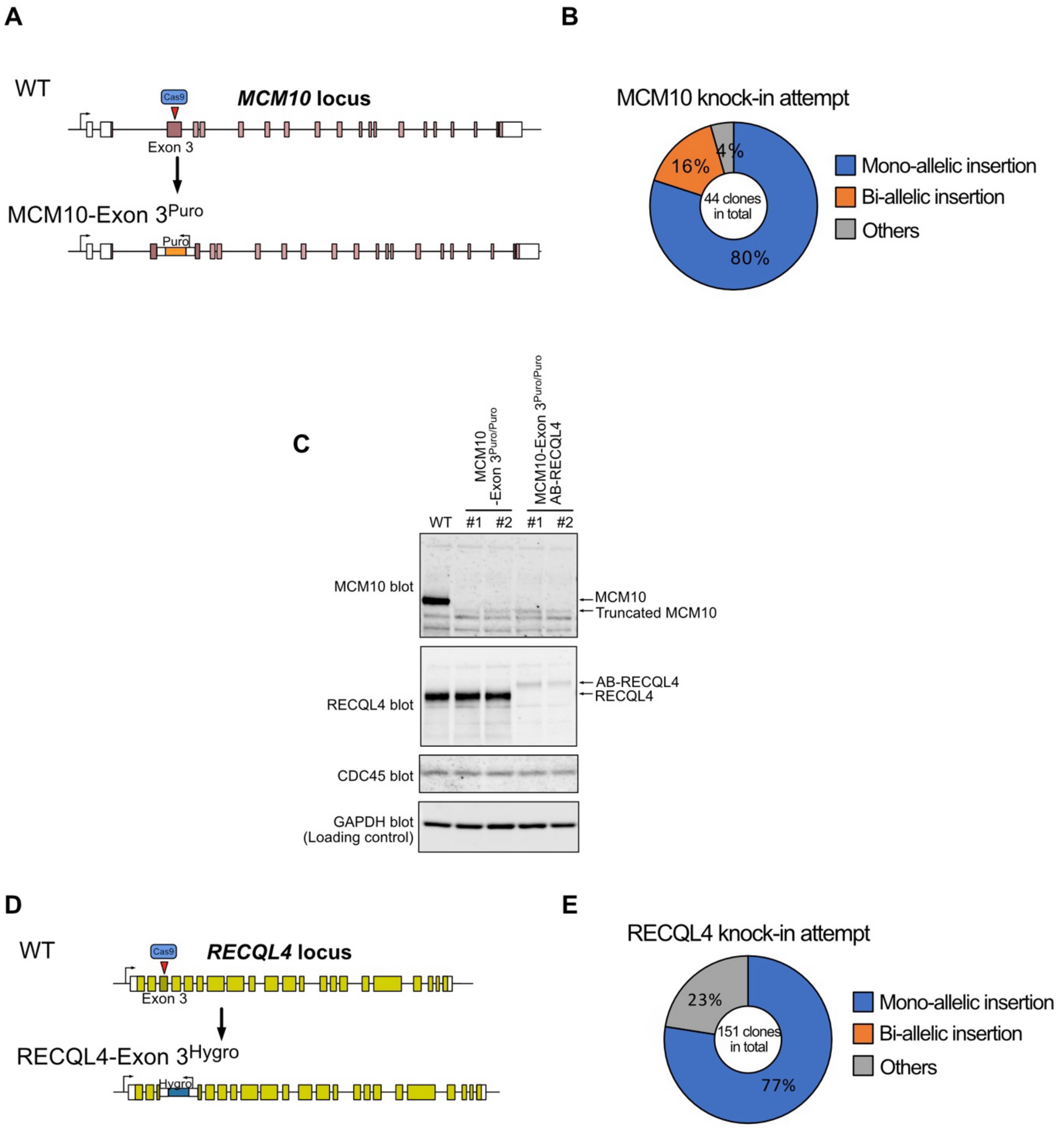


**Figure S1**

**(A)** Schematic showing disruption of exon 3 in the MCM10 gene. MCM10 exons are shown as boxes; uncolored boxes indicate untranslated regions. Exon 3 was a target by CRISPR–Cas9 for insertion of a puromycin-resistance gene cassette. **(B)** Pie chart showing the proportions of mono- and bi-allelic insertion of the puromycin-resistant cassette. **(C)** Immunoblot showing MCM10 expression in the obtained mutants. Two biallelic insertion clones with either wild-type or AID2-BromoTag-RECQL4 background were analysed. All clones exhibited a smaller MCM10 band that was not detected in wild-type cells, suggesting expression of a truncated MCM10 protein. **(D)** Schematic showing disruption of exon 3 in the RECQL4 gene. RECQL4 exons are shown as boxes; uncolored boxes indicate untranslated regions. Exon 3 was targeted by CRISPR–Cas9 for insertion of a puromycin-resistance gene cassette. **(E)** Pie chart showing the proportions of mono- and bi-allelic insertion of the puromycin-resistant cassette.

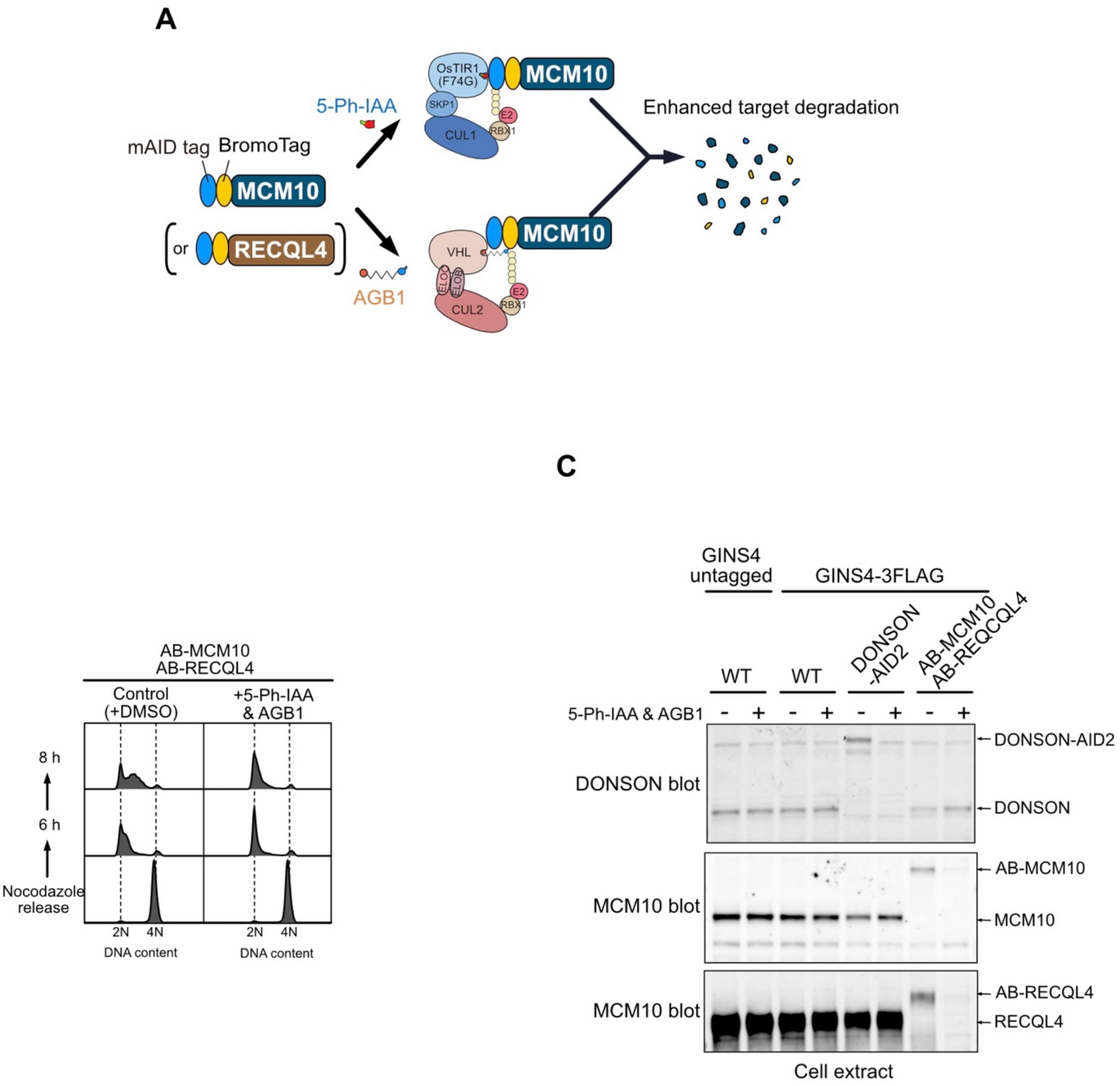


### Figure S2

**(A)** Schematic illustrating how the double-degron system induces protein degradation in the presence of the inducers 5-Ph-IAA and AGB1. **(B)** Cell-cycle progression after nocodazole arrest. The indicated cells were arrested in M phase with nocodazole and then released in the presence or absence of the degradation inducers. DNA content of propidium iodide-stained cells was analysed by flow cytometry. **(C)** Immunoblot showing depletion of the target proteins in the immunoprecipitation assay shown in Fig. 1F. Antibodies used for detection are indicated on the left.

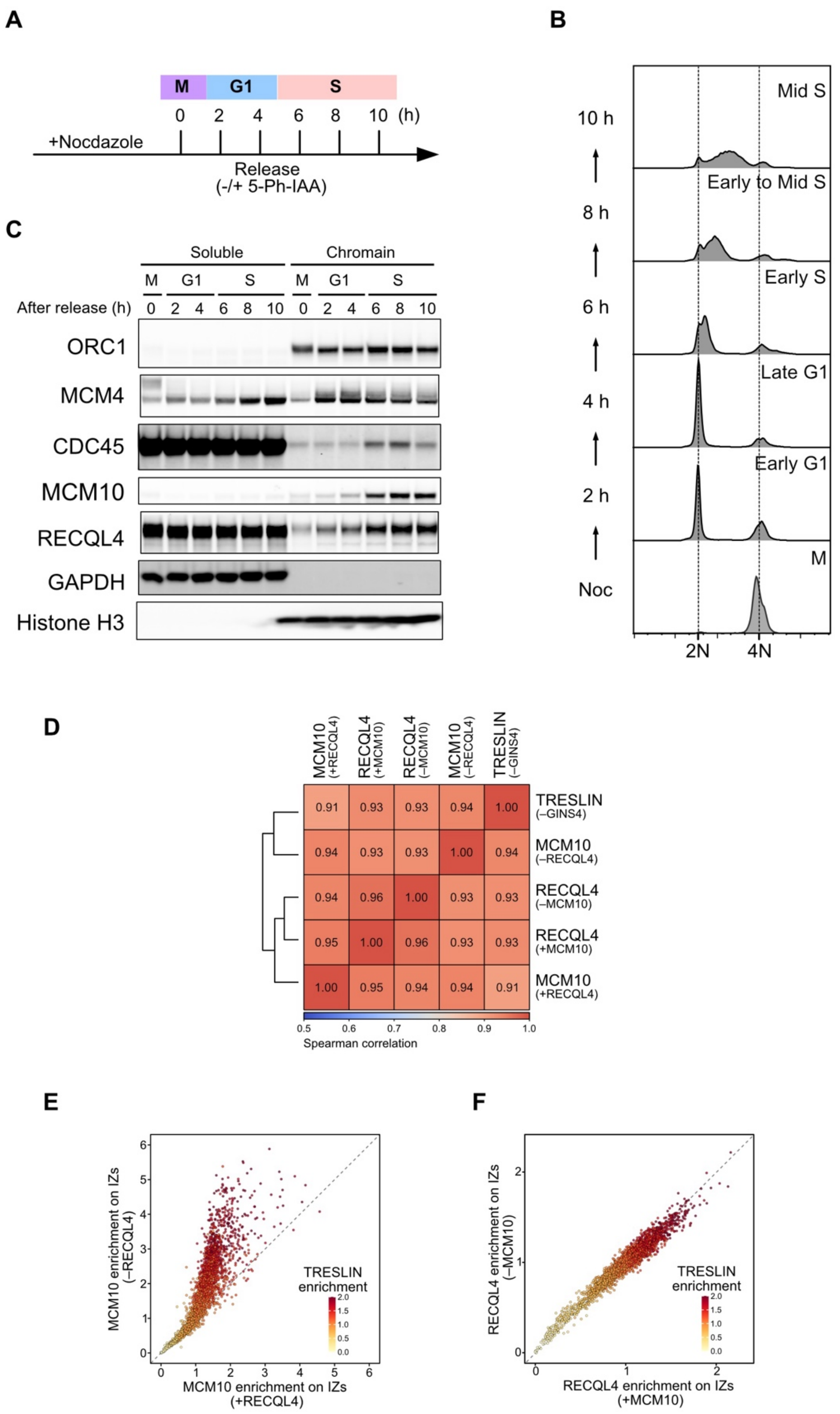


**Figure S3**

**(A)** Schematic illustrating cell-cycle progression after nocodazole release and the time points for sample collection. **(B)** Cell-cycle progression after nocodazole release. DNA content of propidium iodide-stained cells was analysed by flow cytometry. **(C)** Immunoblot showing the timing of chromatin association of the indicated replication factors. Cells collected at the indicated time points after nocodazole release were lysed and then fractionated to obtain soluble and chromatin-bound proteins. **(D)** Spearman correlation matrix showing similarity in enrichment among TRESLIN, MCM10 and RECQL4. Related to Fig. 2D. **(E)** Dot plot showing the accumulation of MCM10 at initiation zones (IZs) in the presence or absence of RECQL4. Red indicates TRESLIN enrichment. **(F)** Dot plot showing the accumulation of RECQL4 at IZs in the presence or absence of MCM10. Red indicates TRESLIN enrichment.

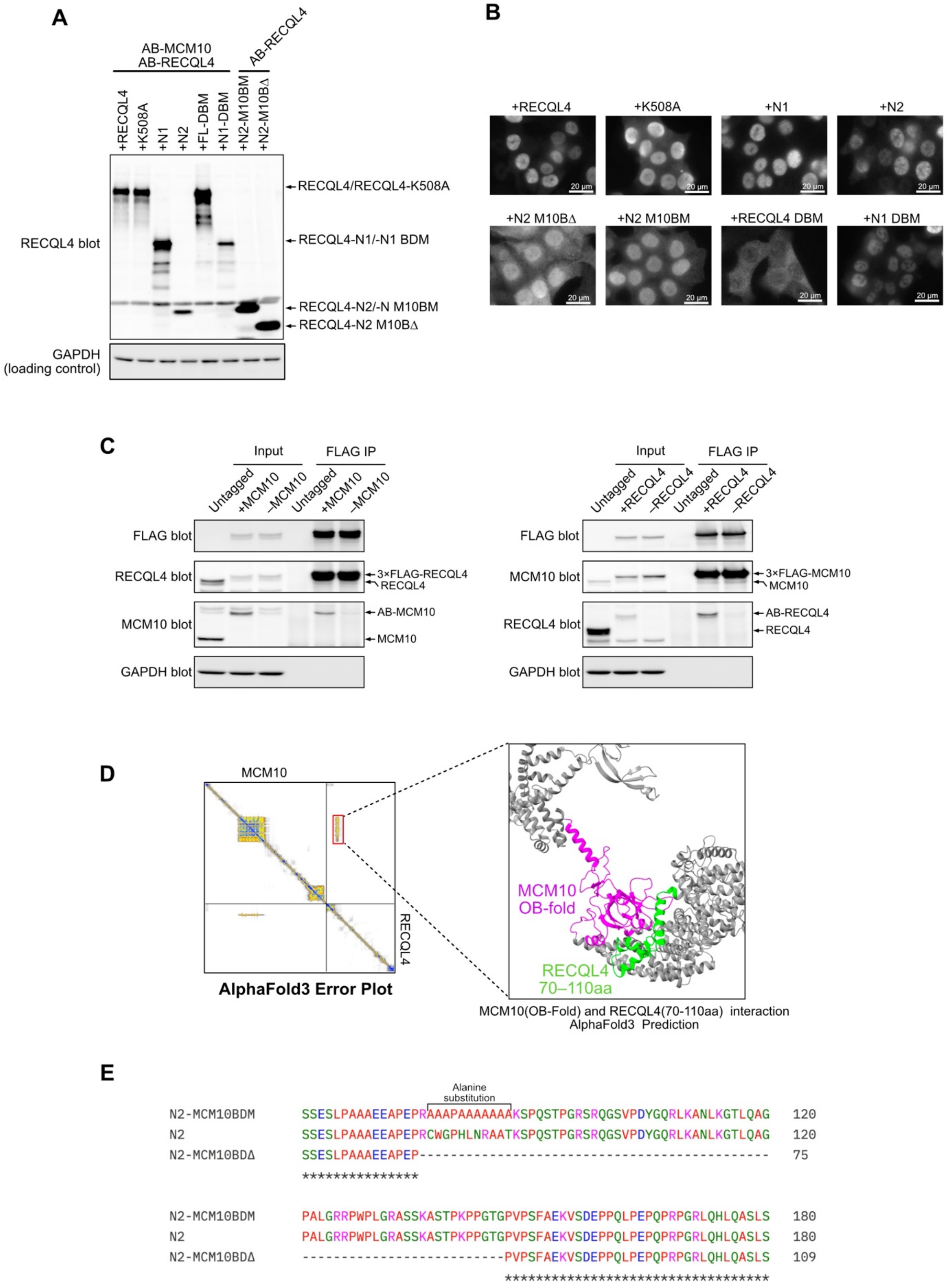


**Figure S4**

**(A)** Immunoblot confirming expression of the RECQL4 mutants used in this study. All proteins were fused to an SV40 nuclear localisation signal to ensure nuclear localisation. **(B)** Cells shown in panel A were stained with an anti-FLAG antibody to confirm nuclear localisation of the ectopically expressed RECQL4 proteins. Cells expressing RECQL4-DBM exhibited both nuclear and cytoplasmic localisation. The scale bar indicates 20 µm. **(C)** Immunoblot showing the interaction between MCM10 and RECQL4. Left, 3×FLAG-RECQL4 was immunoprecipitated in the presence or absence of AB-MCM10. Right, 3×FLAG-MCM10 was immunoprecipitated in the presence or absence of AB-RECQL4. **(D)** AlphaFold3 predicted interaction between MCM10 and RECQL4. Left, predicted error plot. Right, panel highlighting the interacting domains. **(E)** Sequence alignment of RECQL4-N2, -N2 M10DM and -N2 M10DΔ mutants. Substituted and deleted amino acids are indicated.

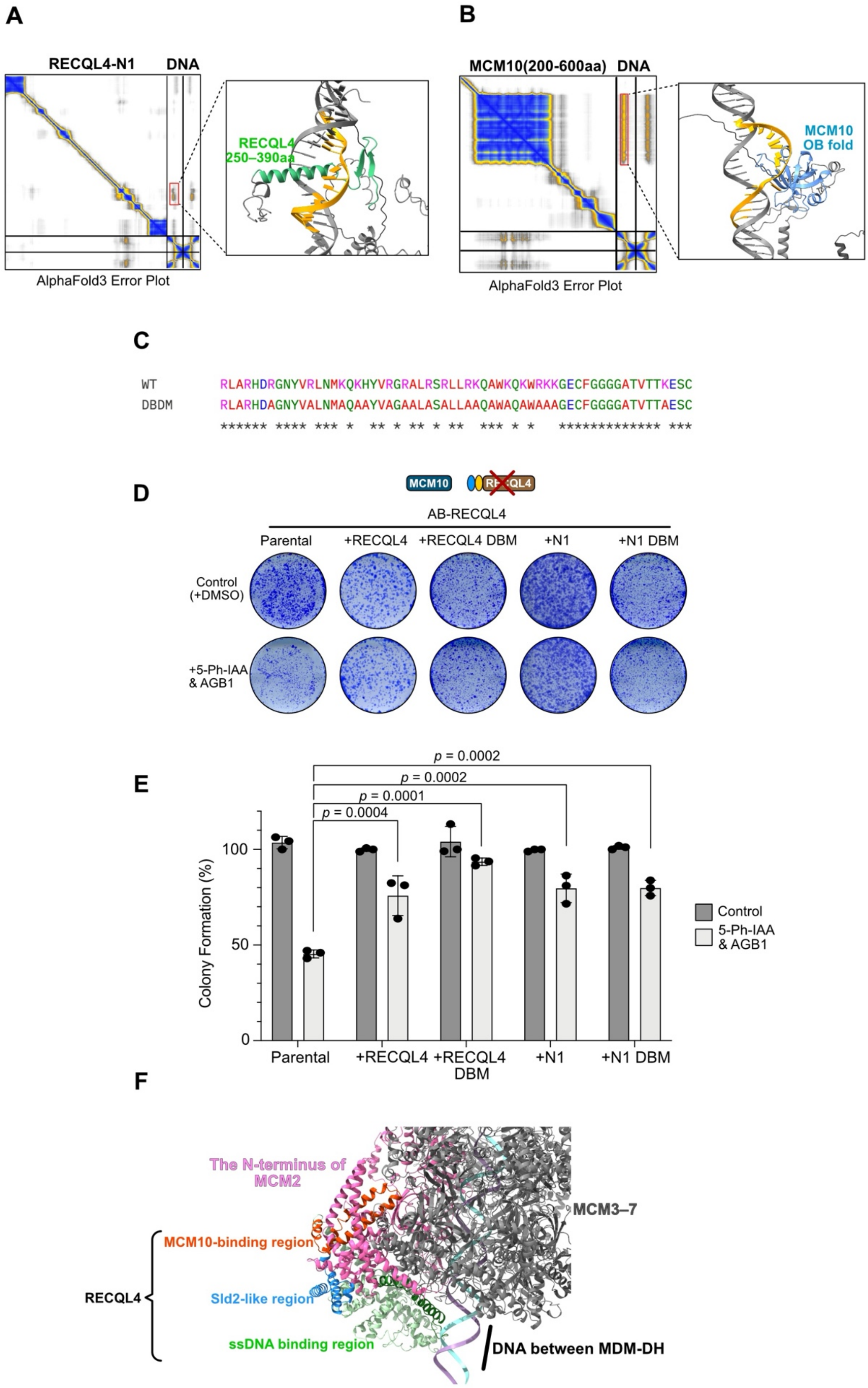


**Figure S5**

**(A)** AlphaFold3-predicted interaction between RECQL4-N1 and bubbled DNA. Left, predicted error plot. Right, panel highlighting the interacting domain and its interaction with ssDNA. **(B)** AlphaFold3 predicted interaction between MCM10 (amino acid 200-600) and bubbled DNA. Left, predicted error plot. Right, panel highlighting the OB-fold domain of MCM10 and its interaction with ssDNA. **(C)** Sequence alignment of RECQL4 and its DNA-binding domain mutant. Substituted amino acids are indicated. **(D)** Representative images of stained colonies. The indicated cells were seeded at 1,000 cells per well and cultured for 7 days in the presence or absence of 5-Ph-IAA and AGB1. **(E)** Quantification of colony formation efficiency in the indicated cells. Data are presented as mean ± SD ($n$ = 3 replicates; one-way ANOVA, multiple comparison) and were normalised to the untreated cells. $p$-values were calculated in comparison to the parental cell line. **(F)** AlphaFold3-predicted interaction between DNA-bound MCM2–7 and RECQL4-N1. The N-terminus of MCM2 is shown in pink. The ssDNA-binding domain of RECQL4 is positioned adjacent to the DNA located between two MCM double hexamers.

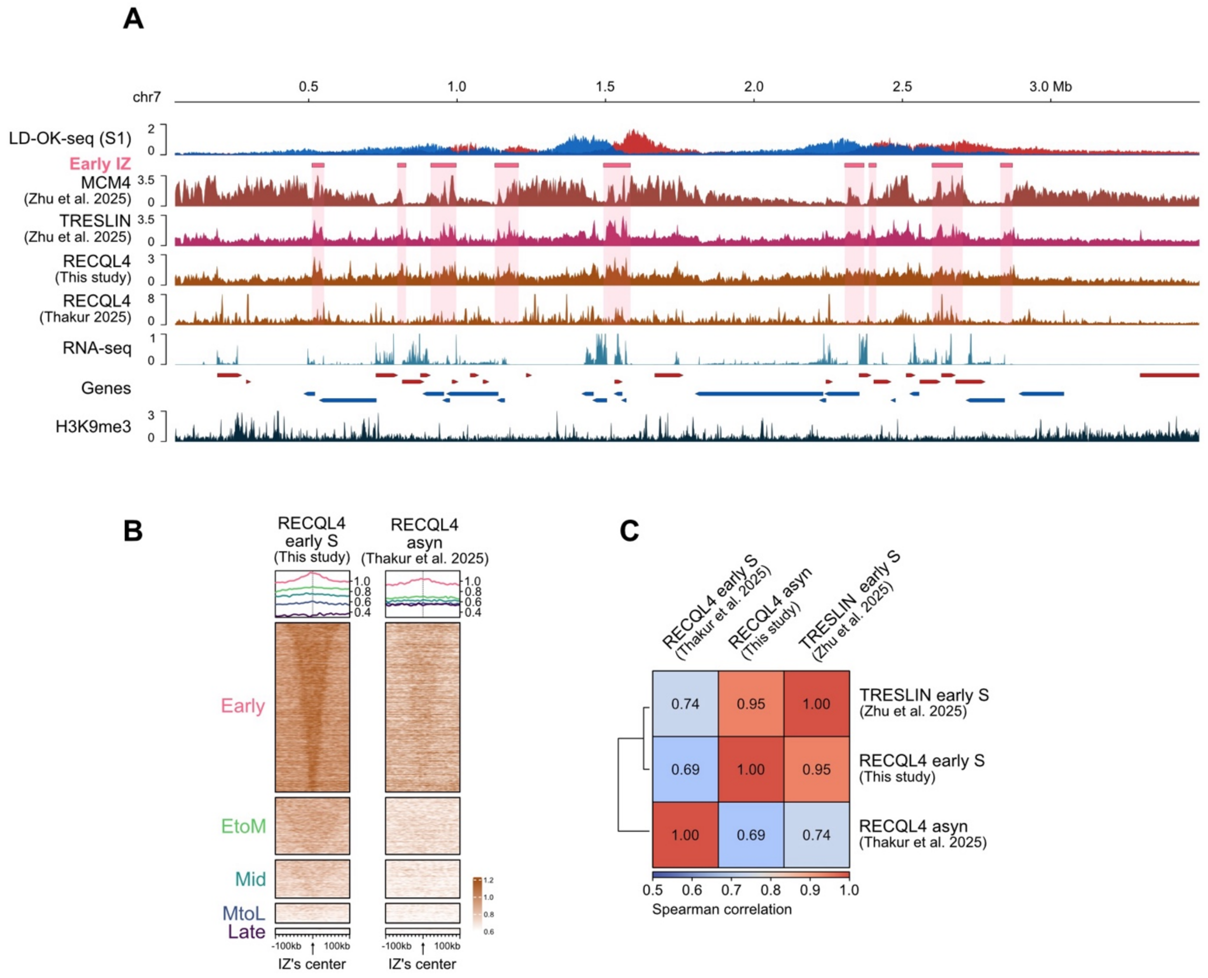


**Figure S6**

Comparison of ChIP-seq profiles of RECQL4 distribution in this study and a recent study. **(A)** Representative genomic profiles of LD-OK-seq in early S-phase (S1), ChIP-seq for MCM4, TRESLIN, RECQL4 (this study and dataset from Thakur et al. 2025), RNA-seq, and H3K9me3 are shown. Early-firing IZs are highlighted in pink. Data from Thakur et al. were analysed using the same pipeline as that used in this study. RNA-seq data are displayed as log2-transformed TPM values. Genes are annotated with coloured arrows indicating transcriptional directions. **(B)** Heatmaps showing RECQL4 enrichment at IZs. RECQL4 ChIP-seq data in this study were obtained from cell in early S phase, whereas the RECQL4 ChIP-seq data reported by Thakur et al. were obtained from asynchronous cells. IZs are categorised according to their activation timing in S phase. IZs are centered with ±100 kb flanking regions, and ordered by size. **(C)** Spearman correlation matrix showing similarity of TRESLIN and RECQL4 ChIP-seq enrichment across the different datasets.
